# Consistency of Sign Language Movement Expression among Proficient and Student Signers

**DOI:** 10.64898/2026.08.17.745267

**Authors:** Eric Harbour, Julia Krebs, Julia Martetschläger, Hermann Schwameder, Dietmar Roehm, Ronnie B. Wilbur, Evie A. Malaia

## Abstract

While movement variability is a natural element of human expression, in sign languages it may affect mutual understanding, learning, and potential overuse injury. Sign language variability is not well-understood in part because quantitative analytical methods are yet to be clearly defined. Hence the aim of this study was to assess intra-subject reliability across repeated sessions for three signers, to identify features sensitive to experience-related differences in motor control consistency, and to establish movement consistency metrics for treating sign language kinematic differences as linguistically meaningful. Three signers were assigned to three different proficiency levels of sign language: Deaf (D), proficient (P), and student (S). Sign production variables were evaluated using intraclass correlation coefficients (ICCs) and coefficients of variation (CVs). Most kinematic features showed good to excellent ICCs such as duration, path length, signing space volume, and average and peak velocity. Some EMG features such as mean forearm amplitudes and co-contraction also showed good to excellent ICCs. These data can be used to improve the scientific investigation of sign languages, improve educational resources, and establish baseline thresholds to inform ergonomic or scheduling guidelines for interpreters.

## 1. Introduction

Movement variability is a natural aspect of behavior that allows our bodies to adapt to fatigue, changing environments, and prevent injury. Variability in sign language movement is influenced by linguistic, sociolinguistic, and individual motor factors, but has broad implications for mutual understanding, learning processes, and also injury risk. About 80% of sign language interpreters report upper extremity repeated stress injuries that have contributed to lapses in their work (Stedt, 1992). Repetitive movement below typical fatigue thresholds (as low as 5% of maximum muscle activation) can still provoke pain via low-threshold motor unit overuse (Roman et al., 2021). Understanding how sign language movement components vary may flag movements prone to overuse injury in interpreters or all-day signers. In addition, quantifying the typical variability of signing may help improve sign language teaching by discerning which aspects differ most between different proficiency levels. Hearing students learning a sign language as a second language (L2) are known to express higher variability than Deaf proficient signers, and the difference narrows with subsequent sign language education (Hilger et al., 2015). Educational resources might be improved with better understanding of which stable factors are associated with proficiency.

3D motion capture is often used to measure small hand movements which are also involved in sign language production. While few studies have directly investigated the specific consistency of sign language movement, studies of other fine and precise non-linguistic movements can provide a relevant comparison. Small hand movements in occupational settings have highly repeatable kinematic features with intraclass correlation coefficient (ICC) values of 0.60 to 0.88 and coefficient of variation (CV) typically from 8.3 to 25.4% (Srinivasan et al., 2015). Violin playing may offer an analogous comparison with its repetitive, fine-motor wrist movements. (Ancillao et al., 2017) reported CVs between 2.7-4.7% for elbow and wrist range of motion (respectively), increasing to 7.2-15.8% for velocity. Furthermore, violin players express average muscle activations around 5-13% maximum voluntary contraction (Mann et al., 2021), which is directly comparable to sign language (Krebs et al., 2025a).

Electromyography (EMG) is an indispensable tool for investigating movement and motor control, with surface EMG especially popular due to its non-invasive and wireless setup. It is commonly used in clinical and scientific settings to differentiate between muscle contraction types and intensities. Forearm EMG shows good consistency within sessions and between days, with ICC values of > 0.8, and reasonable CVs = 5.9% (95% confidence interval = 0.4–12.4) (Sorbie et al., 2018). Generally between-day variation is larger than within-session due to differences in skin preparation and electrode placement, but it is also influenced by fatigue and motivation (Srinivasan et al., 2015).

Within a linguistic system, movement variability is not distributed at random: phonetic-level implementation should remain stable across repetitions (Brentari, 2019; Lucas and Bayley, 2011), while variability that is systematic serves grammatical and/or semantic function (Krebs et al., 2025b; Malaia and Wilbur, 2012; Malaia et al., 2013, 2026). For example, proficient signers have been shown to produce different movement-complexity profiles for verbs with different event structure (low entropy, high predictability for telic verbs; moderate entropy for atelic verbs). This complex motor pattern is acquired well after the gross spatial parameters of a sign have been mastered (Malaia et al., 2026). The asymmetry in acquisition of spatial vs. temporal features of motor control in sign language is similar to that in speech, where spatial articulatory targets (e.g., those responsible for formant frequencies) are approximated early, while temporal targets (e.g., voice onset time) require extended practice and often fail to converge on native norms (Flege, 1995; Flege et al., 1999). Nevertheless, it is still unclear how much variability is in the normal range within-participants, between-days. Understanding which features (kinematic or EMG-based) show stability can clarify skill-related motor patterns. Moreover, cross-sectional studies can show that kinematic and EMG measures differ significantly between conditions (Krebs et al., 2025b), but without an estimate of measurement consistency, an “acute” between-condition difference cannot be distinguished from ordinary within-subject noise. Reliability metrics allow calculation of the minimum detectable change: the smallest between-condition difference that exceeds measurement noise. Without this, single-sign or single-session comparisons risk over-interpreting differences that fall within normal trial-to-trial variability. No study has yet established which kinematic and EMG sign-production features are reliable enough to support this kind of inference in sign language research.

Hence, the objectives of this study were to: 1) assess intra-subject reliability across repeated sessions for three signers, 2) identify features sensitive to experience-related differences in motor control consistency, and 3) establish movement consistency metrics for treating sign language kinematic differences as linguistically meaningful.

## 2. Methods

### 2.1. Participants

Three participants took part in this study, which was a subgroup of a previous analysis (Krebs et al., 2025a). The first signer (D) was a fluent Deaf signer of Austrian Sign Language (ÖGS) who is an active sign language teacher, and is heavily involved in the Deaf community. The second signer (P) was a hearing, highly proficient L2 signer (completed interpreter training; almost 20 years of ÖGS learning/use). The third signer (S) was a hearing L2 sign language student learning ÖGS for 4 months at the time of measurement.

### 2.2. Data Acquisition

Each participant produced a list of 102 signs for two repetitions during a single visit to the laboratory. The list of signs included a mix of verbs (telic and atelic) and adjectives (intensified and non-intensified pairs; i.e., dünn [thin] and sehr dünn [very thin]). The signs were presented visually in written German using a slideshow with one unique slide for each sign. For the student only, videos of the signs were presented to confirm understanding. Participants were able to redo a sign if desired. They were instructed to make a clear separation between reps by starting and stopping each sign with both arms relaxed and extended vertically at their sides. Two of the authors (JK and JM) used the 2D video, 3D recording and wrist trajectory to define the beginning and end of each sign. The sign start and end were identified using a combination of criteria such as handshape, location, and wrist velocity. Motion capture data were gathered using a custom-designed marker set and a 12-camera infrared motion capture system (Qualisys AB, Göteborg, Sweden) operating at a sampling rate of 300 Hz. EMG analysis was performed using EMG sensors (Ultium(TM) EMG, Noraxon, Scottsdale, AZ, USA) connected to surface electrodes (Ambu blue, 30 × 22 mm, Ag/AgCl). Data were collected simultaneously with the kinematic analysis through the Qualisys Track Manager (Qualisys AB, Goteborg, Sweden). EMG data were collected at 2000 Hz. EMG signals were recorded from four arm muscles: m. extensor digitorum, m. flexor digitorum, m. biceps brachii and m. triceps brachii of the dominant arm. EMG electrode placement, and skin preparation was performed according to the best-practice recommendations of SENIAM (http://www.seniam.org/). Participants performed maximum voluntary contraction (MVC) procedures according to best practices (Burden, 2010) by contracting against a fixed object in standardized positions (wrist 0, elbow 90 degrees flexion). They were given strong verbal encouragement to push maximally for five seconds.

### 2.3. Signal Processing

The full processing of the motion capture and EMG data was performed in MATLAB 2026a (The Mathworks, Natick, USA) and was described previously (Krebs et al., 2025a,b). Kinematic data were trimmed to sign duration between the labeled sign start and end. Marker trajectories were filtered with a second-order, zero-lag Butterworth filter with a cutoff frequency of 25 Hz. Then, segment positions and orientations were calculated using an inverse kinematics algorithm (V3D; C-Motion, Rockville, MD, USA). The joint centers of the wrist were estimated as a virtual landmark positioned midway between the lateral and medial anatomical markers; this was the representative point of interest for all subsequent analyses. Linear displacement was calculated as the straight line between the start and end of the sign. Total path displacement was estimated from 3D position data using the cumulative sum of step-wise Euclidean distances between consecutive frames. Three other spatial measures were derived from the 3D wrist trajectory: 3D signing space volume, spatial compactness, and planarity. 3D signing space volume was computed as the volume of the convex hull (MATLAB function “convhull”) enclosing the set of wrist positions, providing an estimate of the overall spatial extent of the sign. Spatial compactness was defined as the convex hull volume of the trajectory normalized by the product of the PCA-derived spatial extents (Quinto et al., 2025):

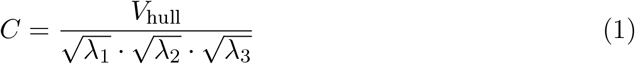

where *V*_hull_ is the volume of the smallest convex shape enclosing all trajectory points, and 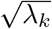 represents the spatial spread along each principal axis. Higher values indicate a more sphere-like, expansive trajectory; lower values indicate a flat or linear one. Planarity was estimated via principal component analysis (PCA) of the *N ×* 3 wrist position point cloud (Ogihara et al., 2014; Promsri and Federolf, 2020). The covariance matrix of the trajectory was decomposed to yield eigenvalues *λ*_1_ ≥*λ*_2_ ≥*λ*_3_, and planarity was defined as the proportion of variance explained by the dominant plane:

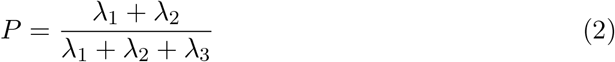

A value of *P* = 1 indicates a perfectly planar trajectory (*λ*_3_ = 0), while lower values reflect increasingly three-dimensional movement. First, second, and third derivatives were calculated using the function “gradient” to obtain velocity, acceleration, and jerk, respectively. Velocity data were transformed using the Euclidean norm to obtain absolute speed, from which the median and peak speed (*m/s*) were labeled. Sample entropy was calculated from the speed vector using the function “SampEn” with an embedded dimension m=2 and tolerance r=0.2*standard deviation (Lee, 2012), and a spatiotemporal index (STI) was calculated using the standard deviation (SD) at every other time point (50 SDs) and then summing all SDs into a scalar value (Howell et al., 2009).

EMG raw data during the sign phase was used to derive the power spectral density (PSD) estimate using the function “periodogram”. Then, mean and median frequency were extracted and the power in five distinct frequency bands was derived for comparison (6-15, 16-25, 26-60, 61-75, and 76-140 Hz) (Roman-Liu and Konarska, 2009). EMG data were further processed with bandpass filtering (10-300 Hz), notch filtering (50 Hz), rectification, and root mean square normalization (100 ms window) according to best practices (Muceli and Merletti, 2024). MVC was extracted from the highest 1s average (Burden, 2010). Sign phase EMG was trimmed and normalized to MVC. The mean, median, and peak (in 0.25 s windows) activation was extracted for comparison. A co-contraction index (CCI) was calculated to approximate the degree of activation between agonist and antagonist muscles in the dominant arm (e.g., upper arm biceps and triceps) (Li et al., 2021)). The formulas of (Rudolph et al., 2000) and (Falconer, 1985) were adapted such that each index was quantified at each point to determine the higher- and lower-amplitude signal independently at every time point rather than fixing a single agonist for the entire trial. This was done since certain sign expressions had alternating agonist muscles within the sign. At each sample *i*, let EMG_low_(*i*) and EMG_high_(*i*) denote the lower- and higher-amplitude signal, respectively, of the two antagonist channels. The continuous co-contraction indices were calculated as:

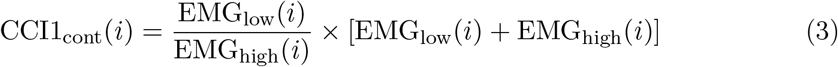

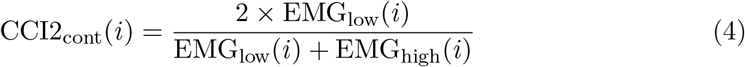

CCI1 (Rudolph formulation) increases with both the relative and absolute magnitude of co-activation, whereas CCI2 (Falconer–Winter formulation) is a normalized ratio bounded between 0 and 1, reflecting only the relative balance between the two signals. Mean and peak (0.25 s window) CCI during the sign phase were retained for comparison.

### 2.4. Statistical analysis

The consistency of sign production variables was evaluated using accepted reliability metrics intraclass coefficients (ICC) and coefficient of variation (CV). An ICC(3,1) model of absolute agreement with two-way mixed-effects was used to determine intra-subject consistency. Intraclass correlation coefficients were interpreted using the conventional benchmarks of poor (<0.50), moderate (0.50–0.75), good (0.75–0.90), and excellent (>0.90) reliability (Koo and Li, 2016). CV was computed per sign as the quotient of the within-sign standard deviation across reps and the within-sign mean × 100. CVs were averaged across all signs. ICCs and CVs were calculated separately per participant (D, P, S) and per feature, using rep 1–rep 2 paired signs. Typical error is presented instead of CV for time to peak deceleration, planarity, and CCI2 to accommodate the finite limits of those variables. Wilcoxon signed-rank tests were used to explore the presence of absolute differences in key metrics between participants. Reps were collapsed into a single scalar value and then compared pairwise within sign.

## 3. Results

Across most kinematic and EMG features, the Deaf (D) and proficient (P) signer showed comparable good-to-excellent ICCs, while the student was more variable. Kinematic variables generally showed good to excellent ICC (many >0.85) with CVs commonly 10–20%. EMG variables were more mixed, with PSD variables showing the greatest variability.

### 3.1. Kinematics

All three signers were generally consistent in their expression of wrist displacement and velocity (Table 1). Total path length and straight displacement ICCs ranged from 0.93–0.98 (Figure 1). Peak wrist speed also had excellent ICC (0.90–0.94) with CV tightly clustered at 10–11%. Peak deceleration also had good consistency with ICC = 0.81–0.88 and CV = 17-18%. Finally, STI wrist velocity was also good to excellent (ICC = 0.88–0.96).

**Table 1.**
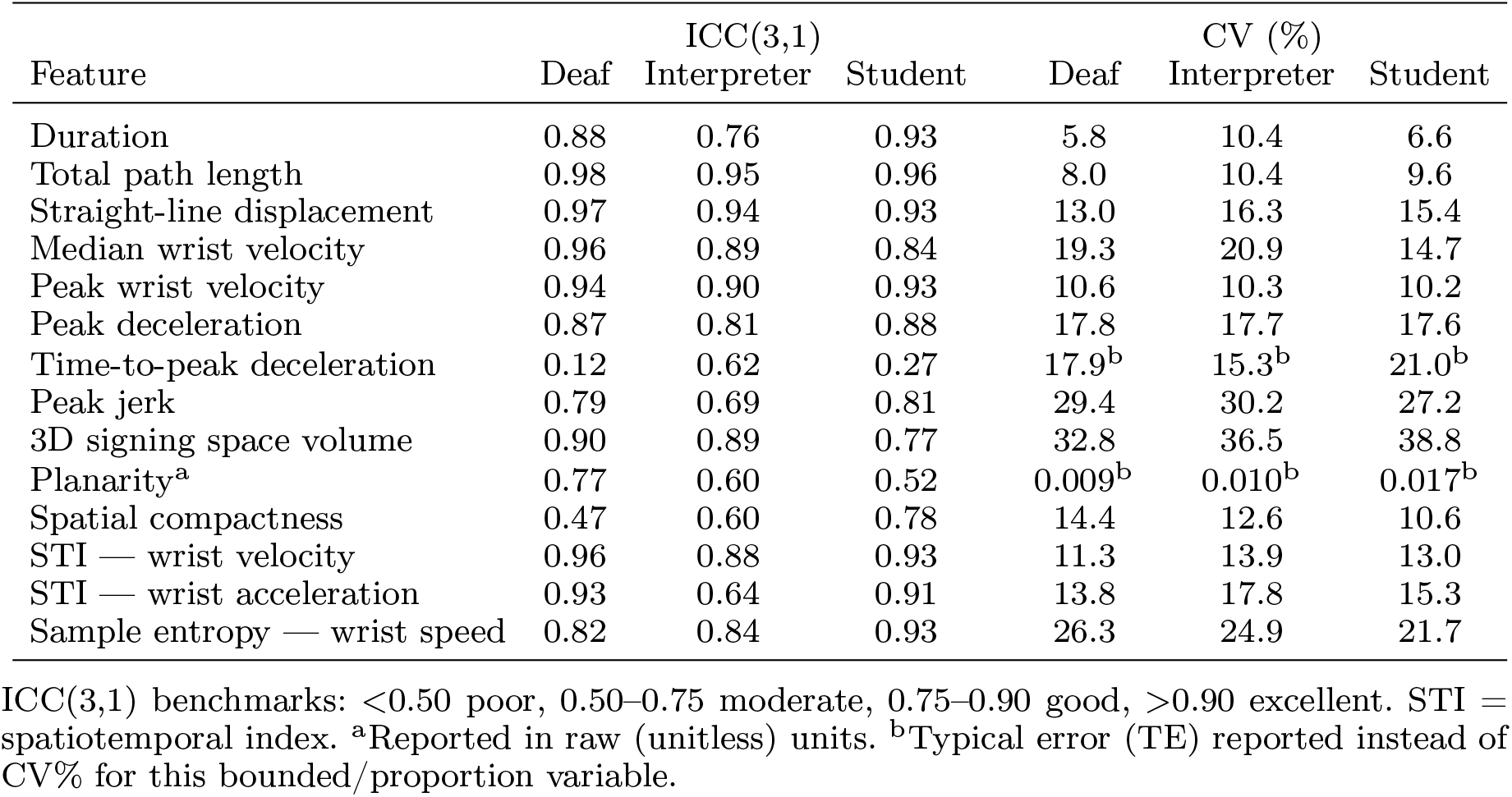
Intra-subject reliability (ICC(3,1), CV%, and TE) for spatiotemporal/kinematic features by signer.

**Figure 1.**
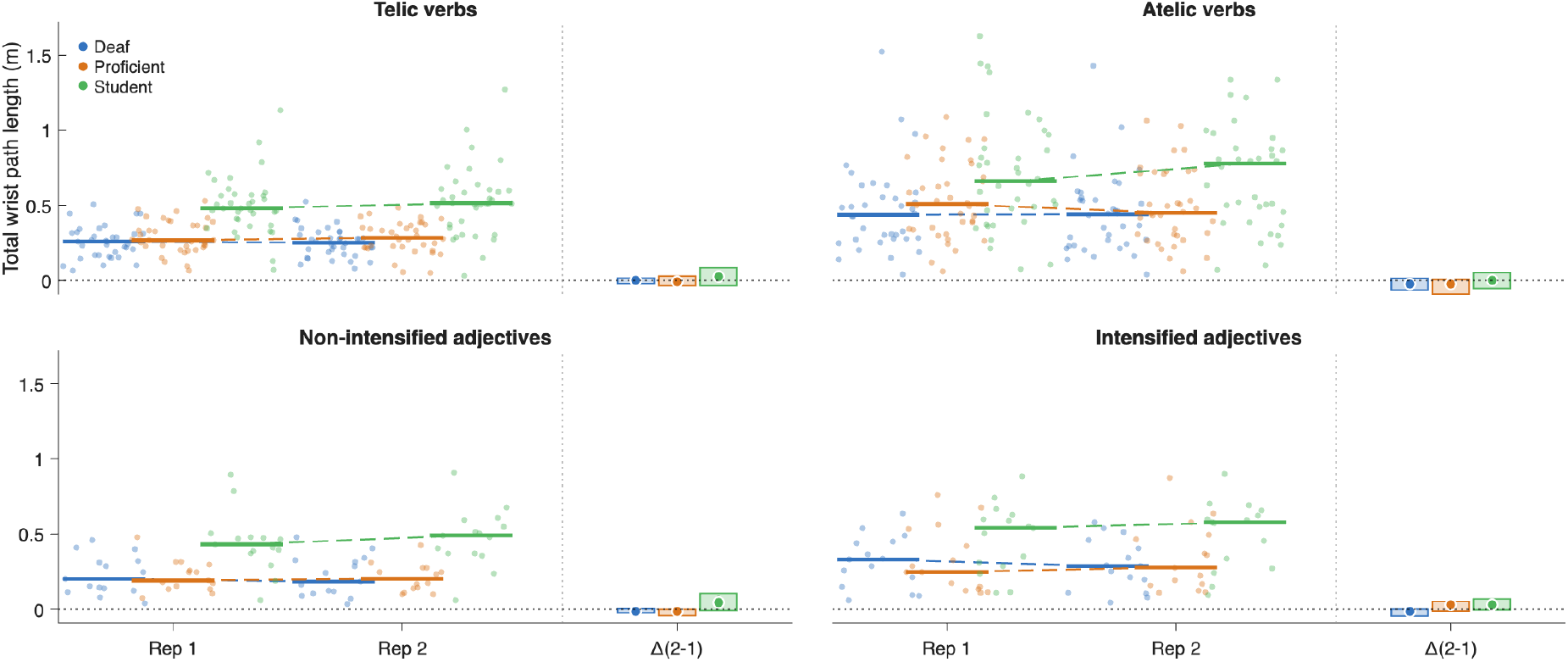
Gardner-Altman style difference plots of total wrist path length. Note the average higher path length for the student, while the Deaf signer and proficient are similar. Each dot represents the discrete value for one sign repetition (rep), with distinct colors for each participant. Median values are visualized with solid horizontal bars, with the dashed line connector representing the median within-subject difference between repetitions. The panel adjacent to rep 2 displays the between-rep median difference (dot), inter-quartile range (line), and 95% confidence interval (shaded box).

Some metrics showed marked variation between signers. Signing space volume was consistent for D and P (ICC = 0.89–0.90), but less so for the student (S; ICC = 0.77). While planarity showed good reliability for D (ICC = 0.77), it was moderate for P (0.60) and poor for S (0.52). STI acceleration had excellent ICCs for D and S (0.91–0.93) but was only moderate for P (0.64). Unexpectedly, S showed the highest ICC values in the group for sign duration (0.93), spatial compactness (0.78), and wrist speed sample entropy (0.93) (Figure 2). Time to peak deceleration consistency was uniformly poor across signers (ICC = 0.12–0.62, typical error = 15.3-21.0%).

**Figure 2.**
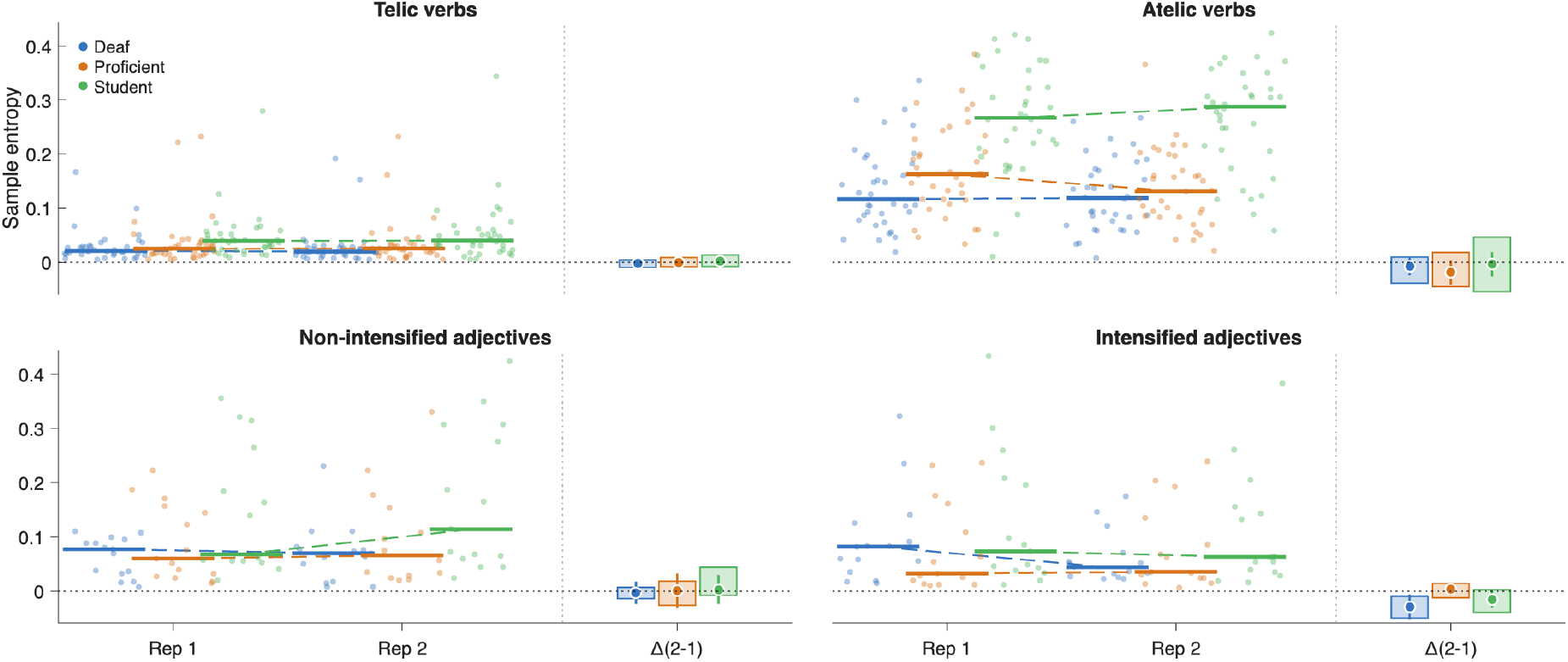
Gardner-Altman style difference plots of wrist speed sample entropy. Note the higher absolute values for the student signer in atelic verbs.

### 3.2. EMG

EMG measures displayed substantially more variability than kinematic variables (Table 2). Average forearm amplitudes had mostly good to excellent ICCs ranging 0.70–0.92 and CVs from 10-18% (Figure 3). Upper arm EMG ICCs ranked mostly good as did peak EMG across all muscles. Moreover, EMG activation consistency mostly followed D > P > S. The same pattern was apparent in the flexor PSD features, where D’s ICCs (0.80–0.93) consistently exceeded P and S whose were moderate to poor. For the triceps EMG amplitudes S reliability generally matched or exceeded D and P (e.g., mean triceps EMG ICC = 0.91 S vs. 0.86 for D and 0.74 P). Average PSD values also displayed inconsistent and paradoxical reliability values (e.g., extensors mean frequency ICC = 0.97 for S > 0.74 for P > 0.41 for D).

**Table 2.** Intra-subject reliability (ICC(3,1) and CV%) for EMG amplitude, frequency, and co-contraction features by signer.

| Feature | ICC(3,1) |  |  | CV (%) |  |  |
| --- | --- | --- | --- | --- | --- | --- |
|  | Deaf | Interpreter | Student | Deaf | Interpreter | Student |
| *Flexors* |  |  |  |  |  |  |
| Mean amplitude | 0.89 | 0.88 | 0.76 | 10.3 | 17.1 | 11.3 |
| Median amplitude | 0.78 | 0.83 | 0.70 | 11.6 | 17.1 | 12.6 |
| Peak amplitude | 0.93 | 0.80 | 0.69 | 13.4 | 24.2 | 14.9 |
| MPF | 0.84 | 0.64 | 0.77 | 7.6 | 6.2 | 6.1 |
| MF | 0.80 | 0.53 | 0.69 | 11.4 | 8.8 | 8.7 |
| PSD bands<sup>a</sup> | 0.86±0.03 | 0.60±0.15 | 0.66±0.10 | 33±8 | 44±7 | 36±11 |
| *Extensors* |  |  |  |  |  |  |
| Mean amplitude | 0.91 | 0.89 | 0.81 | 15.9 | 13.3 | 18.0 |
| Median amplitude | 0.92 | 0.90 | 0.81 | 15.5 | 13.2 | 18.1 |
| Peak amplitude | 0.75 | 0.83 | 0.70 | 21.7 | 17.2 | 22.8 |
| MPF | 0.60 | 0.85 | 0.98 | 10.6 | 4.5 | 9.7 |
| MF | 0.41 | 0.74 | 0.97 | 15.5 | 6.8 | 9.1 |
| PSD bands<sup>a</sup> | 0.70±0.14 | 0.67±0.19 | 0.66±0.22 | 43±13 | 35±9 | 38±7 |
| *Biceps* |  |  |  |  |  |  |
| Mean amplitude | 0.70 | 0.87 | 0.70 | 29.9 | 14.8 | 14.3 |
| Median amplitude | 0.74 | 0.84 | 0.77 | 25.4 | 16.4 | 13.0 |
| Peak amplitude | 0.03 | 0.84 | 0.47 | 45.1 | 16.8 | 21.1 |
| MPF | 0.31 | 0.62 | 0.61 | 9.4 | 8.0 | 6.5 |
| MF | 0.26 | 0.61 | 0.48 | 11.7 | 11.2 | 9.4 |
| PSD bands<sup>a</sup> | 0.50±0.19 | 0.73±0.10 | 0.53±0.08 | 60±10 | 40±11 | 36±9 |
| *Triceps* |  |  |  |  |  |  |
| Mean amplitude | 0.86 | 0.74 | 0.91 | 19.8 | 17.7 | 10.3 |
| Median amplitude | 0.75 | 0.67 | 0.84 | 18.3 | 16.7 | 11.3 |
| Peak amplitude | 0.83 | 0.54 | 0.83 | 29.6 | 22.8 | 15.7 |
| MPF | 0.70 | 0.79 | 0.56 | 11.7 | 6.6 | 6.1 |
| MF | 0.57 | 0.56 | 0.43 | 15.8 | 10.0 | 7.7 |
| PSD bands<sup>a</sup> | 0.73±0.14 | 0.28±0.22 | 0.69±0.20 | 46±16 | 43±10 | 32±12 |
| *Co-contraction indices* |  |  |  |  |  |  |
| CCI1 — forearm | 0.89 | 0.81 | 0.83 | 11.0 | 19.7 | 12.2 |
| CCI2 — forearm | 0.87 | 0.88 | 0.73 | 0.032<sup>b</sup> | 0.066<sup>b</sup> | 0.061<sup>b</sup> |
| Peak CCI1 — forearm | 0.93 | 0.76 | 0.74 | 15.0 | 27.1 | 15.5 |
| Peak CCI2 — forearm | 0.72 | 0.86 | 0.50 | 0.068<sup>b</sup> | 0.093<sup>b</sup> | 0.056<sup>b</sup> |
| CCI1 — upper arm | 0.84 | 0.82 | 0.85 | 19.0 | 18.6 | 11.2 |
| CCI2 — upper arm | 0.80 | 0.84 | 0.74 | 0.055<sup>b</sup> | 0.068<sup>b</sup> | 0.064<sup>b</sup> |
| Peak CCI1 — upper arm | 0.75 | 0.82 | 0.64 | 26.2 | 24.2 | 17.0 |
| Peak CCI2 — upper arm | 0.69 | 0.77 | 0.65 | 0.079<sup>b</sup> | 0.105<sup>b</sup> | 0.076<sup>b</sup> |

*Table 2 continued*
| Feature | ICC(3,1) |  |  | CV (%) |  |  |
| --- | --- | --- | --- | --- | --- | --- |
|  | Deaf | Interpreter | Student | Deaf | Interpreter | Student |
| ICC(3,1) benchmarks: <0.50 poor, 0.50–0.75 moderate, 0.75–0.90 good, >0.90 excellent. <sup>a</sup>Power spectral density collapsed across the seven frequency bands per muscle; values are mean ± SD of the per-band ICC/CV. <sup>b</sup>Typical error (TE) reported instead of CV% for CCI2 (Falconer–Winter formulation), which is bounded 0–1 and for which CV is not well defined; CCI1 (Rudolph formulation) is not bounded on a comparable scale and CV is retained. CCI = co-contraction index; MPF = mean power frequency; MF = median frequency, PSD = power spectral density. |  |  |  |  |  |  |

**Figure 3.**
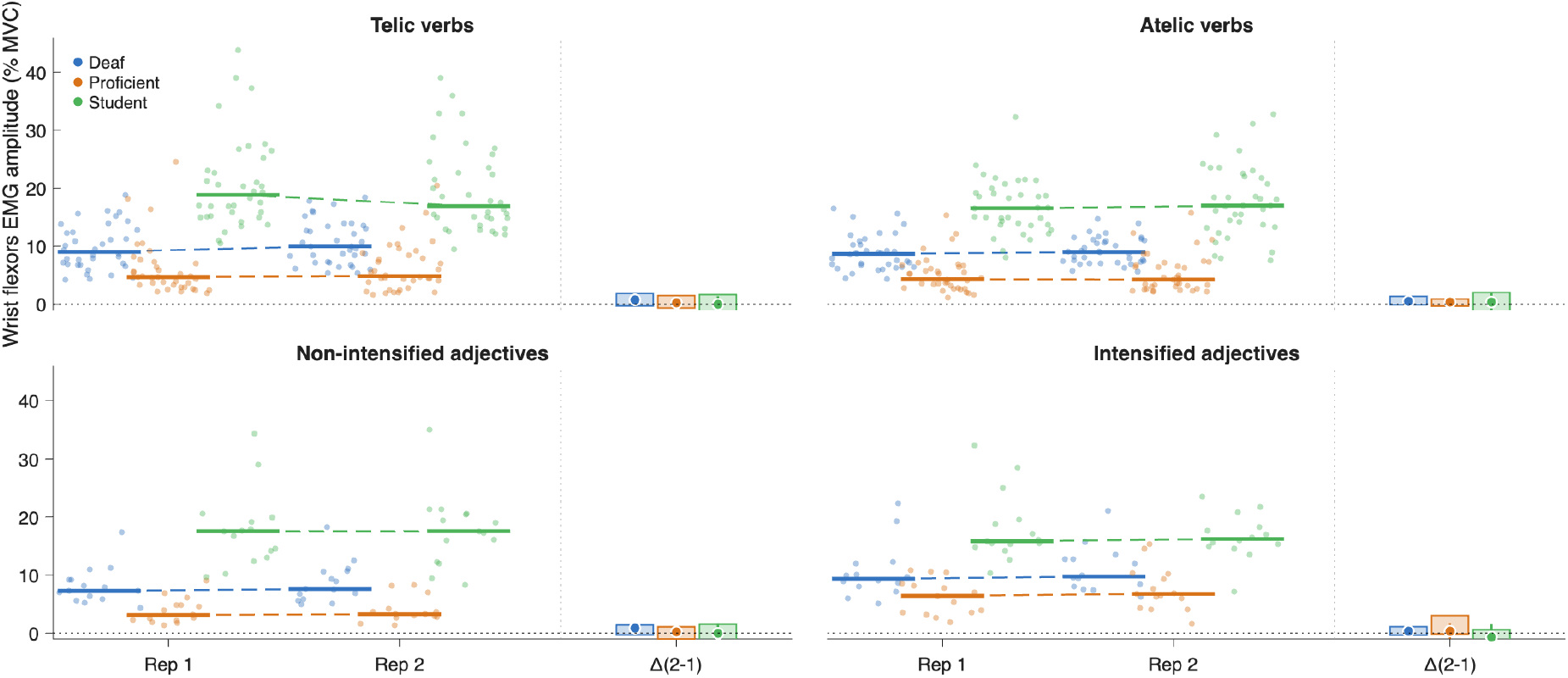
Gardner-Altman style difference plots of mean wrist flexor EMG amplitude. Note the average higher activation for the student across all sign types, despite stable values between repetitions.

Co-contraction indices showed good reliability, with average co-contraction ICCs = 0.73–0.89 and CV = 7–20%. While peak values had slightly lower consistency, all indices showed similar ICC and CV values between signers.

## 4. Discussion

This is the first study of its kind reporting the consistency of sign language biomechanics. These findings suggest that most spatiotemporal and kinematic features are stable across proficient and student signers. Muscle activation was more variable, but amplitude features (scaled to MVC) showed variability comparable to the former, with many “good” ICC values. Hence, between-subject differences in absolute values (i.e., in path length, Figure 1) likely reflect real differences in movement expression and support previous observations of differences in kinematics and EMG between word types (Krebs et al., 2025a).

### 4.1. Intra-subject Consistency

Many features were highly stable among all three signers (good to excellent ICCs), including duration, path length, signing space volume, average and peak velocity, peak deceleration, and sample entropy. EMG features such as mean forearm amplitudes and CCI1 also showed good to excellent ICCs. These metrics can be used with sufficient confidence in scientific investigation to compare proficiency levels, word types, or perhaps to evaluate the effects of fatigue. Other variables did not demonstrate meaningful consistency: time to peak deceleration, upper arm EMG amplitudes and most PSD features exhibited poor-to-moderate reliability across the sample. These metrics should be avoided or used with caution since the intra-subject variability was high. Other studies examining the consistency of small upper-extremity movements have shown that within usually exceeds that of between-subjects kinematic (Srinivasan et al., 2015) and EMG (Oskouei et al., 2013) variability. The features that showed the highest reliability (i.e. duration, path length, and displacement) are the gross spatiotemporal parameters that L2 signers acquire within weeks of instruction, well before they gain control over fine-grained velocity-profile structure (Malaia et al., 2026). On the other hand, the features that showed the weakest reliability (i.e. time-to-peak deceleration, planarity) depend on the trajectory of articulation across the full time course rather than its gross endpoints, and are also the last kinematic properties to stabilize even after months of instruction (Malaia et al., 2026).

### 4.2. Unifying and distinguishing features

All three participants showed not only low variability but also similar absolute values for peak wrist deceleration (-0.00011-0.00013 m/s^2^; Figure 4) and peak jerk (0.000029-0.000037 m/s^3^). These kinematic features might be especially relevant for between-word comparisons or injury risk study designs. While only the Deaf signer showed “good” ICC, all had identical planarity (0.99); hence, this metric may have limited specificity or sensitivity in sign language research.

**Figure 4.**
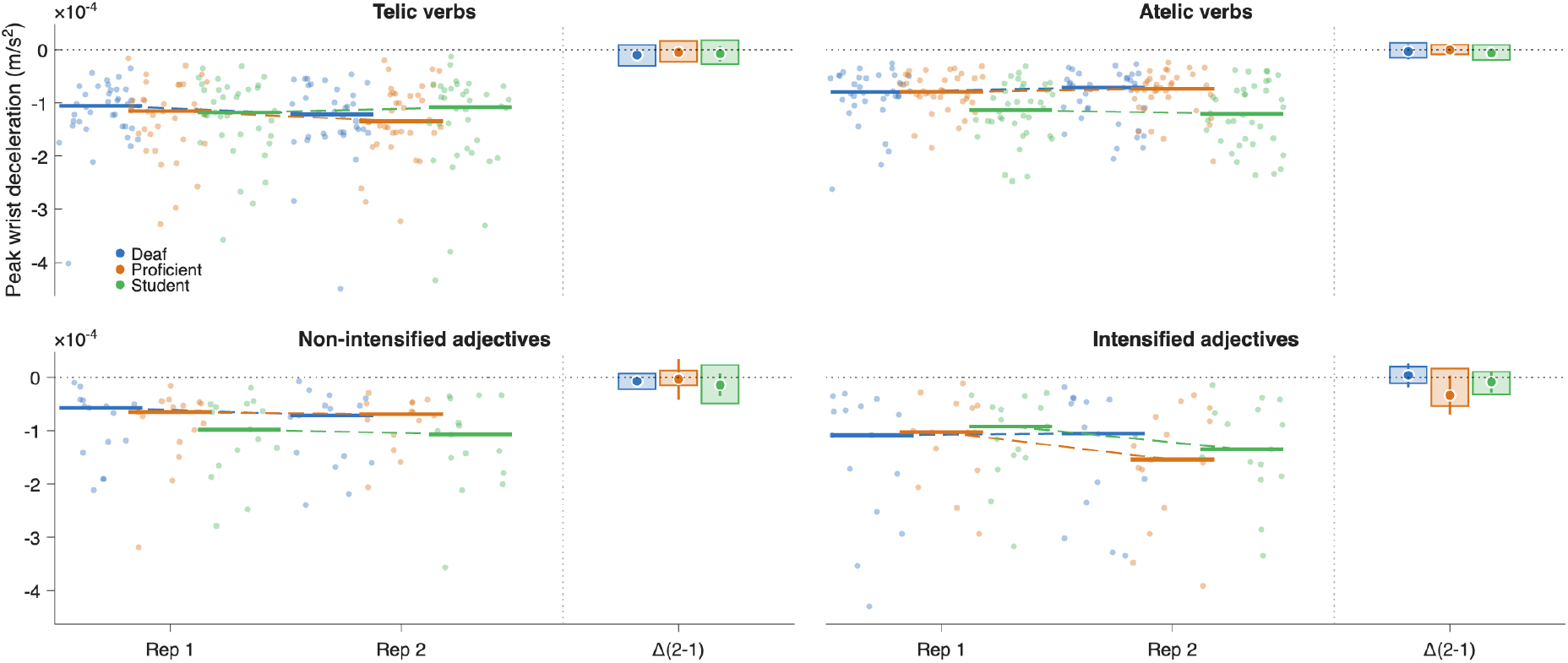
Gardner-Altman style difference plots of peak wrist deceleration. Note the similar deceleration values between subjects and repetitions within each sign type.

The convergence across signers at all proficiency levels on peak deceleration and jerk, despite otherwise substantial between-signer differences in absolute magnitude, is consistent with these features functioning as *phonetic* motor targets (Blumenthal-Dramé and Malaia, 2019; Torre et al., 2019). The reliability profile reported here, with some features stable across all signers, others reliably differentiating experience level, and some unstable regardless of proficiency, is consistent with sign production kinematics being linguistically organized rather than just reflecting motor control variability.

Due to the relatively stable spatiotemporal and kinematic expression within signers, the current sample showed statistically meaningful differences in several parameters. The student (S) differed from the Deaf (D) and proficient (P) signers in straight displacement (0.25 vs. 0.14-0.15 m, p < 0.05) and STI wrist velocity (0.0085 vs. 0.0057-0.0059 m/s, p < 0.05). Notably, P and S diverged from D in almost all parameters such as path length (D = 0.33, P = 0.37, and S = 0.57 m, all p < 0.05), peak velocity (0.0026, 0.0030, and 0.0038 m/s) and CCI2 of the forearm (0.18, 0.31, and 0.70, respectively, all p < 0.05). This observation indicates that there are inherent aspects of proficiency unique to Deaf signers that distinguish them from highly competent but hearing interpreters. This mirrors previous studies that found that increased sign language education reduces variability (measured by STI) but even high competency does not converge with the movement variability of Deaf signers (Hilger et al., 2015).

While there are no exact comparisons for these findings in sign language research, observations in upper-extremity precision work or music are likely relevant since repetitive, fine-motor control is connected to practice, proficiency, and overuse injuries. In the current study, the average within-participant ICCs for kinematic features was 0.74-0.76, which is exactly in the middle of observed values in occupational settings (Srinivasan et al., 2015). However, the CVs reported here (18-21%) are high in the range of (Srinivasan et al., 2015) (8 to 25%) and much higher than those seen in violin players (7-16%) (Ancillao et al., 2017). To contextualize the EMG findings is even more difficult with few relevant comparisons; two studies examined within-day EMG repeatability of a various handgrip tasks and reported good to excellent ICCs (> 0.8) (Oskouei et al., 2013) and CVs of 5.9% (95% CI = 0.4–12.4) (Sorbie et al., 2018). The current findings reflect lower consistency, with ICCs ranging 0.58-0.76 and CVs of 25-33%. This could be attributed to the lower absolute muscle activation and dynamic nature of sign language, both of which likely interact to reduce observed measurement consistency. Hence, a portion of the observed variability is due to natural intra-subject differences in EMG activation; any measurement error could be further reduced by using improved EMG systems incorporating electrode arrays or intramuscular electrodes.

### 4.3. Limitations

Due to the resource-intensive nature of 3D motion capture, this study included a limited number of participants; hence, these findings should be considered primarily exploratory. Other Deaf signers serving as a “reference” might express different absolute values and consistency because of base language personal signing style. While the single-day, cross-sectional design may have limited representation of each signer’s consistency, inter-day study designs were avoided due to confounding effects of day-to-day fatigue, electrode placement, and EMG normalization (Oskouei et al., 2013; Sorbie et al., 2018). Finally, the Deaf signer’s observed biceps EMG amplitudes were probably vulnerable to the latter; activation levels of 133-218% MVC are not physiologically feasible. Since both reps were similarly mis-scaled, this error did not likely affect the measured reliability values.

### 4.4. Conclusion

This study is one of the first to report the variability and consistency of meaningful biomechanical features of sign language across different proficiency levels. Most kinematic features had high consistency, which emphasizes not only their specificity but also sensitivity to changes in proficiency, linguistic meaning, or injury. Muscle activation (when scaled to MVC) also showed good consistency within-subjects. These data can be used to improve the scientific investigation of sign language, improve educational resources, and establish baseline thresholds to inform ergonomic or scheduling guidelines for interpreters. Future work should deploy the most stable metrics to larger multi-signer datasets, and use them to discover the variability and generalizability of movement patterns. Feature selection or dimensionality reduction could be used to better classify or automate proficiency, for example in educational resources.

## Acknowledgement(s)

The researchers extend their sincerest gratitude to the study participants who generously volunteered their time in the laboratory.

## Funding

This research was funded in whole or in part by the Austrian Science Fund (FWF) [10.55776/ESP252 and 10.55776/P35671], and by the National Science Foundation [grant numbers 1932547, 1734938].

## Notes on contributor(s)

EH - Writing - Original Draft, Formal analysis, Investigation, Methodology, Visualization, Data Curation; JK - Writing - Review & Editing, Conceptualization, Funding Acquisition, Investigation, Project Administration; JM - Writing - Review & Editing, Data Curation; EM - Writing - Review & Editing, Conceptualization, Funding Acquisition, Supervision; RBW - Writing - Review & Editing, Conceptualization, Funding Acquisition, Supervision; HS - Writing - Review & Editing, Funding Acquisition, Resources; DR - Writing - Review & Editing, Funding Acquisition.

## Notes

### Competing Interest Statement

The authors have declared no competing interest.

